# Translation elongation factor EF-P drives susceptibility to fluoroquinolone antibiotics in *Bacillus subtilis*

**DOI:** 10.64898/2026.09.27.754791

**Authors:** Madison Ahmad, Rodney Tollerson, Kevin S. Lang

## Abstract

The emergence of antimicrobial resistance restricts the clinical efficacy of therapeutic antibiotic drugs. Elucidating the molecular mechanisms by which bacteria evade antibiotic activities will lead to novel therapeutic approaches. Therefore, it is critical to understand the mechanisms that govern susceptibility to antibiotics. Fluoroquinolones are a class of antibiotics widely used to treat both Gram-positive and Gram-negative bacterial infections. Resistance to fluoroquinolones is widespread in multiple pathogenic bacteria, limiting their therapeutic usefulness. Resistance to fluoroquinolones is well-known to be mediated by Point mutations in the genes encoding the cellular targets of fluoroquinolones, type-II topoisomerases, leads to clinically relevant resistance. However, several recent studies have revealed that dysregulation of translation influences fitness during fluoroquinolone exposure. For example, several screens have identified insertions in gene that encodes elongation factor P (EF-P) increases survival to fluoroquinolones. Here, we sought to understand the impact of EF-P on fluoroquinolone survival using the Gram-positive model organism *Bacillus subtilis.* We found that loss of EF-P increases the survival of *B. subtilis* exposed to fluoroquinolones by several orders of magnitude. Furthermore, we demonstrate that fluoroquinolone sensitivity is dependent on post-translational modification of EF-P, indicating that it is dependent on EF-P activity. Remarkably, loss of EF-P increases survival of cells lacking RecA, a key DNA repair protein, which is crucial for the survival of fluoroquinolone drugs. Using transcriptomics, we identify key cellular response pathways differentially regulated in cells lacking EF-P during fluoroquinolone treatment. We conclude that EF-P regulates diverse mechanisms that drive the susceptibility to fluoroquinolones.

## INTRODUCTION

Antimicrobials are crucial for fighting bacterial infections. Antimicrobial resistance (AMR) is eroding the utility of frontline antibiotics, making it essential to understand how bacterial cells respond to drug-induced damage. Recent analyses have estimated that AMR bacteria will be responsible for more human deaths than cancer by 2050 [1]. Beyond treating bacterial infections, antibiotics are also important protecting human health during a multitude of other clinical interventions including chemotherapy, organ transplant, intensive care, and routine surgical procedures [2]. As rates of AMR increase, these interventions become progressively riskier, as common infections can no longer be reliably prevented or treated. The develop novel effective antimicrobial therapies, it is necessary to better understand how bacterial cells respond to drug treatments. While the principal molecular targets of most antibiotics are well described, the physiological responses and how they drive cell death remain unclear [3–5].

Fluoroquinolone antibiotics target the Type II topoisomerases DNA gyrase and Topoisomerase IV [6–8]. Intuitively, mutations in genes encoding Type II topoisomerases confer resistance to fluoroquinolones [9]. These drugs trap topoisomerases as they cleave DNA, resulting double-stranded DNA breaks (DSBs) and chromosomal fragmentation. Fluoroquinolone-induced DSBs result in activation of the bacterial DNA damage response pathway called the SOS system [10, 11]. Cells deficient in DSB repair are hypersensitive to fluoroquinolone treatment, likely due to the necessity of repairing chromosomes fragmented after removal of poisoned topoisomerases on DNA [12–15]. Recent genome-wide analyses of transposon mutants have revealed that the cellular response to fluoroquinolone treatment is more complex than previously thought. For example, genome-wide screens from multiple organisms have shown that disruption of translation regulation confers increased fitness to fluoroquinolone exposure [12, 16–19]. However, the role of translation in regulating fluoroquinolone susceptibility is not well known.

Elongation Factor P (EF-P) is a universally conserved translation elongation factor, orthologous to archaeal aIF5A and eukaryotic eIF5A, that alleviates ribosome stalling at polyproline motifs [20, 21]. EF-P binds near peptidyl-transferase center and facilitates peptide bond formation to restore nascent peptide elongation. In its absence, ribosomes pause during translation of two or more consecutive proline residues because proline is the slowest peptide bond donor and acceptor [22]. Though EF-P is conserved, in bacteria loss of EF-P confers diverse phenotypes. For example, EF-P is essential in *Neisseria gonorrhea,* while in *Bacillus subtilis* loss of EF-P has no impact on vegetative growth [23, 24]. Intriguingly, Tn-Seq experiments in *Salmonella typhi* have suggested that loss of EF-P results in increased fitness during fluoroquinolone treatment. Recent ribosomal profiling studies in *B. subtilis* have shown that EF-P regulated proteins are functionally enriched for DNA replication and repair proteins [25]. While these findings suggest a role for EF-P in regulating the response to DNA damaging drugs, such as fluoroquinolones, such a function has yet to be documented.

Here, we use the Gram-positive model organism *B. subtilis* to investigate the role of EF-P in response to fluoroquinolone treatment. We found that loss of EF-P significantly increased survival to ciprofloxacin, a widely used fluoroquinolone antibiotic. This effect is dependent on post-translational modification of EF-P by YmfI. Additionally, our results suggest that expression of the critical DNA repair protein, RecA, is reduced in cells lacking EF-P, likely due to a conserved polyproline motif. However, in cells lacking RecA, loss of EF-P confers increased survival to ciprofloxacin exposure. RNA-seq experiments show upregulation of genes involved in envelope remodeling, teichoic acid modification, and osmotic protection in cells lacking EF-P during ciprofloxacin treatment, along with changes in redox metabolism and reduced induction of ROS-generating pathways. Together, our results suggest that EF-P shapes survival of fluoroquinolone treatment by through direct or indirect control of both cell-envelope and oxidative stress pathways.

## METHODS

### Growth conditions

*Bacillus* strains were grown in Lysogeny Broth (LB) (10g tryptone, 5g yeast extract, 5g NaCl per liter) or on LB agar (1.5% agar) plates. Ciprofloxacin (Sigma Aldrich) was added to the media at the indicated concentrations. When appropriate, antibiotics to select for gene knockouts were used at the following concentrations: 7µg/mL kanamycin (kan). All incubations were performed at 37° C. Liquid cultures were incubated with shaking at 260 RPM.

### Strain construction

All gene knockouts used in this study were acquired from the Bacillus Genetic Stock Center (https://bgsc-osu.nbsstore.net/). All knockouts or complementation constructs were transformed into *B. subtilis* JH642 (KL1) by natural transformation as previously described [26]. The bacterial strains used in this work are listed in Table S1. All primers used in this study are listed in Table S2.

### Plasmid construction

To create an EF-P complementation construct, we used Gibson cloning [27]. First, plasmid pBSE3Klux was amplified with primers KL323 and KL324. The Pveg promoter was amplified from KL1 genomic DNA using primers KL325 and KL326. The *lacZ* gene was amplified from plasmid HM181 with primer KL327 and KL328. All products were amplified using Q5 PCR master mix (NEB), checked on an agarose gel, and purified using a Thermo GeneJet PCR purification kit. Equal molar amounts of PCR products were mixed in a standard Gibson reaction and transformed into *E. coli* DH10B cells using a standard heat shock method. Next, P*veg*-*lacZ* was amplified from the resulting plasmid using primers KL401 and KL402. The *lacA* integration backbone was amplified from plasmid pBS2EPxylA_V2 using primers KL399 and KL400. These two products were ligated together using a standard Gibson reaction. Finally, the *lacA* integration vector containing the P*veg* promoter was amplified from the resulting plasmid using primers KL583 and Kl584. Primers KL557 and KL558 were used to amplify *efp* with its endogenous RBS. These two fragments were cloned together using a standard Gibson reaction to make the plasmid pKL48.

### Survival assays

Strains were grown by inoculation of a single colony from agar plates into 2 mL LB broth and incubated with shaking at 37° C to exponential phase. They were then normalized to OD600 0.5 and subsequently serially diluted using 10-fold dilutions in phosphate buffered saline (PBS). A 5 µL aliquot of each dilution was plated onto LB agar plates with and without the indicated concentrations of ciprofloxacin. Plates were then incubated at 37° C before colony counting and imaging. Percent survival was calculated as the ratio of colony forming units/mL (CFU/mL) on plates containing drug compared to the CFU/mL on plates without drug.

### GFP quantification assays

Strains were grown by inoculation of a single colony from agar plates into 2 mL LB broth and incubated with shaking at 37° C to exponential phase. They were then normalized to OD600 0.5 in a black-walled, clear bottom 96-well plate (Costar) and the experimental conditions were treated with ciprofloxacin. The plates were incubated for 30 minutes at 37° C prior to reading them on a 96 well plate reader (BioTek). For each well, the 488 nm wavelength intensity was measured, as well as the OD600. GFP fluorescence was normalized to cell density by dividing these two values.

### RNA-Seq

Strains were grown in triplicate by inoculation of a single colony from agar plates into 2 mL LB broth and incubated with shaking at 37° C to an OD600 of ∼1.0. Flasks containing 5 mL of LB broth were inoculated to an OD600 of 0.05 and grown for 1 hour. Ciprofloxacin was added to half the cultures at a final concentration of 50 ng/mL for 30 minutes. Cultures were then dumped into an equal volume of ice cold 100% methanol (Sigma Aldrich) to fix the cells. Cells were pelleted by centrifugation and the methanol was removed. Total RNA was extracted using a GeneJet RNA purification kit (Thermo). A total of 10 µg of total RNA from each sample was treated with DNAse I (NEB) for 30 minutes at 37° C. After heat inactivation, RNA was purified using a GeneJet RNA purification Kit (Thermo). Purified RNA samples were sent on dry ice for ribosomal RNA depletion and next generation sequencing using Illumina paired-end (2×150bp) library preparation at SeqCoast Genomics.

### RNA-Seq Analysis

Resulting FASTQ files were subjected to quality control using FastQC 0.12.1. Low quality bases and adapters were trimmed using Trimmomatic 0.39. Removal of residual rRNA was done using SortMeRNA 4.3.7. Psuedoalignment and read counts per gene were done using Salmon 1.10.3. Normalization and differential expression analysis was conducted using DEseq2. Gene-set enrichment analysis was done using fGSEA 1.28.0.

### Pulse-field gel electrophoresis (PFGE)

Chromosomal DNA was prepared in agarose plugs to minimize mechanical shearing. Cultures were grown in 10mL of fresh LB medium and at 37 °C with shaking to mid-exponential phase (OD₆₀₀ ≈ 0.3–0.4). Cells were treated with 50ug/mL of ciprofloxacin. Untreated and treated samples were taken after 20 minutes of growth and killed with sodium azide to a concentration of 0.02%. Wash out samples were pelleted and washed three times in PBS, then released into LB to grow for 30 and 60 minutes. Cell pellets were collected by centrifugation at 5000 RPM for 3 min at 4°C for all samples. Cell pellets were transferred to Eppendorf tubes and cell density was adjusted to 0.3 OD600. Molten 1.6% low–melting point agarose (Bio-Rad) equilibrated to 55 °C were mixed gently with lysozyme and the cell pellet and immediately dispensed into Bio-Rad plug molds. Plugs were allowed to solidify at 4 °C for 30-40 min.

Agarose plugs were transferred to lysis buffer containing 0.5 M EDTA (pH 8.0), 1% N-lauroyl sarcosine, and lysozyme (final concentration 1 mg/mL) and incubated at 37°C for 5h to digest the *B. subtilis* cell wall. Plugs were then incubated overnight at 50°C in proteinase K buffer (0.5 M EDTA, 1% N-lauroyl sarcosine, 0.5 mg/mL proteinase K) to lyse cells and remove protein–DNA complexes. Following lysis, plugs were washed three times in TE buffer at room temperature to remove residual detergent and proteinase K. Plugs were stored in TE buffer at 4°C until electrophoresis.

Chromosomal DNA was separated using pulsed-field gel electrophoresis in a CHEF-DR III system (Bio-Rad). Agarose plugs were loaded into a 1% pulsed-field certified agarose gel prepared in 0.5X TBE buffer. Electrophoresis was performed at 14 °C in 0.5× TBE under the following conditions: Voltage gradient: 6 V/cm, Switch time: initial 50s, final 90s, Run time: 20h, included angle: 120°. Following electrophoresis, gels were stained in 1% SYBR Gold (Invitrogen) for 30 minutes and imaged using a gel documentation system under UV illumination. Intact chromosomal DNA migrated as a high–molecular-weight band, whereas fragmented DNA appeared as diffuse signal migrating below the chromosome band. Chromosome fragmentation was quantified by measuring the intensity of the intact chromosome band using ImageJ (NIH).

## RESULTS

### Loss or inactivation of EF-P increases survival to ciprofloxacin

EF-P has been shown to regulate translational stalling of hundreds of proteins that contain polyproline motifs in *B. subtilis* [25]. Analysis of these proteins shows that DNA replication and repair proteins are functionally enriched, suggesting that EF-P is involved in regulating the response to DNA damage [25]. However, Tn-Seq studies in *S. typhi* have suggested that loss of EF-P results in increased fitness during treatment with ciprofloxacin, a DNA damaging drug. To address this discrepancy, we measured the survival of wild-type (WT) and Δ*efp* cells during exposure to ciprofloxacin. We found that cells lacking EF-P were less susceptible to ciprofloxacin treatment by several orders of magnitude (Fig. 1). This phenotype was restored by complementation when *efp* was expressed from a constitutive promoter at an ectopic site on the chromosome (Fig. 1). Importantly, this phenotype is not due to slower growth, as we found that cells lacking EF-P grow at similar rates to WT cells (Fig. S1). We assessed whether this effect was specific to ciprofloxacin or if cells lacking EF-P exhibit increased survival to a broad spectrum of antibiotics. We tested the survival of WT and Δ*efp* cells to rifampicin (a transcription inhibitor) and gentamicin (a translation inhibitor). We found that survival of these drug treatments was no different between the two strains (Fig. S2A). Ciprofloxacin, like other fluoroquinolone drugs, induces DSBs [8]. When we tested survival to other DNA damaging agents, we found that loss of EF-P increases to survival to other potent DSB-inducing compounds (mitomycin C (MMC) and phleomycin), but not DNA base damaging agents (4-Nitroquinoline 1-oxide (4NQO) and Methyl methanesulfonate (MMS)) (Fig. S2B-E). We conclude that sensitivity to ciprofloxacin, MMC, and phleomycin is dependent on EF-P.

**Figure 1.**
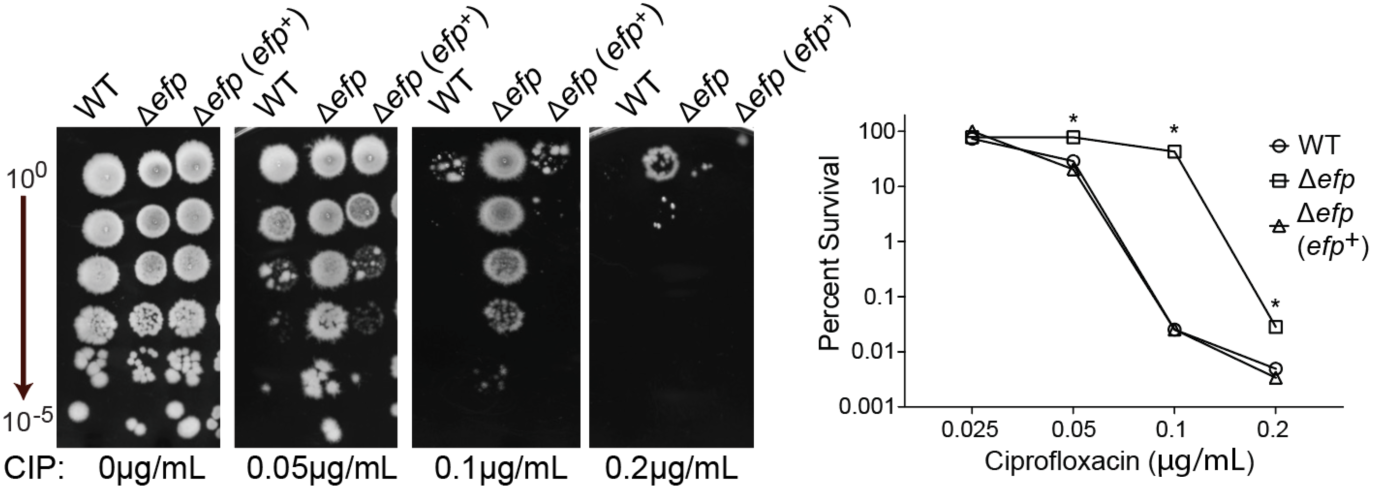
Loss of EF-P increases survival to ciprofloxacin. Representative plates and quantification of ciprofloxacin survival assays. Percent survival was calculated as colony forming units (CFU) on ciprofloxacin plates vs no drug controls. *Efp^+^* indicates cells harboring an *efp* complementation construct. Data represent the mean ± SEM from biological replicate experiments (n=3). * Indicates P-value <0.05.

EF-P is universally conserved across all organisms. In many organisms that have been experimentally characterized, EF-P requires post-translational modification (PTM) to function. The structure of these modifications and the enzymes that ligate them differ across species [24, 28, 29]. In *B. subtilis*, the EF-P PTM is 5-aminopentanol at a conserved lysine residue at amino acid position 32 [30]. Previous studies have suggested that several enzymes are required to assemble the 5-aminopentanol PTM in a stepwise manner, rather than a single ligation step and that each stepwise modification intermediate has a different impact on EF-P activity [30]. The final step of the pathway relies on Ymfl to ligate 5-aminopentanol to lysine-32. We tested whether these PTMs impacted survival to ciprofloxacin. In the absence of drug, all strains grew indistinguishably from WT, indicating that the deletions did not cause a general growth defect, consistent with previous studies. Following ciprofloxacin exposure, a YmfI deletion mutant phenocopied the *efp* mutant, suggesting that active rescue of stalled ribosomes by EF-P drives susceptibility to ciprofloxacin (Fig. 2) [30]. We included a Δ*yfmR* mutant in these experiments because YmfR has recently been shown to reduce translational stalling at polyproline motifs in *B. subtilis* [31]. We found that cells lacking YmfR have no change in susceptibility of ciprofloxacin compared to WT, likely because it is proposed to function when EF-P is absent. We conclude that sensitivity of WT cells to ciprofloxacin requires a 5-aminopentanol PTM on EF-P and that intermediate EF-P PTM moieties do not impact survival.

**Figure 2.**
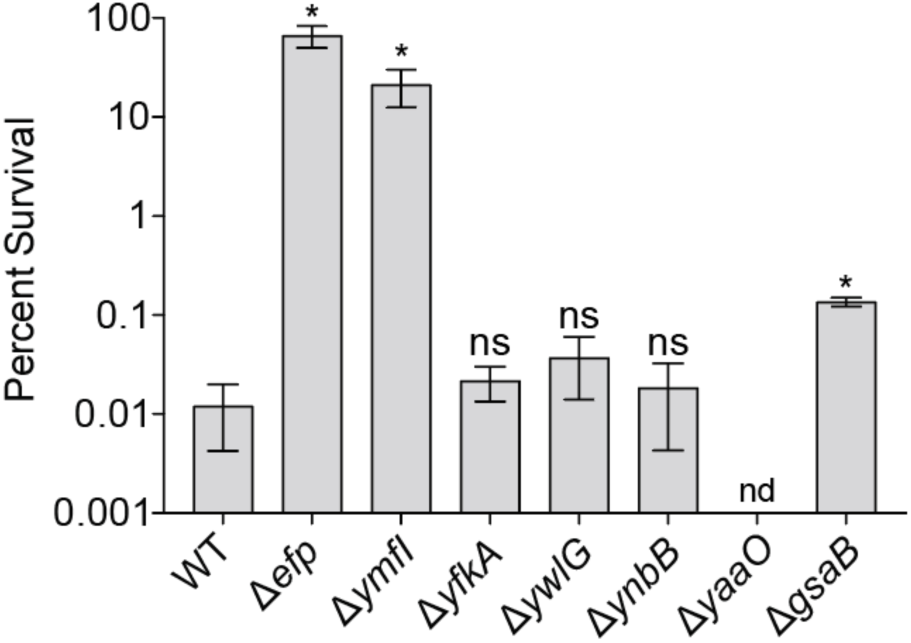
Loss of EF-P post translational modification increases survival to ciprofloxacin. Quantification of ciprofloxacin survival assays. Percent survival was calculated as colony forming units (CFU) on ciprofloxacin plates (50ng/mL) vs no drug controls. Data represent the mean ± SEM from biological replicate experiments (n=3). * Indicates p-value <0.05. ns indicates not statistically significant.

### Increased survival of Δ*efp* cells during ciprofloxacin treatment is due to functions upstream of DNA repair

Previous studies in *B. subtilis* have shown that EF-P-dependent ribosome stalling is enriched at transcripts encoding DNA replication and repair proteins [25]. RecA is a critical DNA repair factor responsible for processing DNA breaks, regulating the SOS response, and survival of DNA damage [32, 33]. *B. subtilis* RecA contains a polyproline motif that is conserved across diverse bacterial species (Fig. 3A). Given the critical role of RecA in regulating the SOS response, we investigated whether EF-P regulates the production of RecA. We monitored the production of a RecA-GFP fusion protein before and after ciprofloxacin treatment in WT and Δ*efp* cells [34]. In untreated cells, RecA-GFP production was reduced in cells lacking EF-P compared to WT (Fig. 3B). As expected, after ciprofloxacin treatment, RecA-GFP levels increased in a dose-dependent manner (Fig. 3B). In cells lacking EF-P, RecA-GFP levels increased after ciprofloxacin treatment, but were reduced significantly compared to WT cells (Fig. 3B). These results are consistent with previous experiments in *B. subtilis* that demonstrate that while RecA is required for DNA damage survival, SOS induction is not [35]. We conclude that RecA production is dependent on EF-P. We infer that this effect is due to the polyproline motif in RecA. Given these results, we considered the possibility that the increased survival of *efp* mutants to ciprofloxacin is due to a mechanism upstream of repair of DNA breaks. To determine this, we measured the survival of Δ*recA* and Δ*recA* Δ*efp* cells during ciprofloxacin treatment. We found that inactivation of *recA* resulted in hyper-sensitivity to ciprofloxacin, as we had to perform these experiments a lower concentration of drug, suggesting that DNA repair is still required for survival of *efp* mutants. However, Δ*recA* Δ*efp* cells were less sensitive compared to Δ*recA* single mutants (Fig. 3C). Additionally, we analyzed genome fragmentation by pulse field gel electrophoresis and found that Δ*efp* cells experience less genome fragmentation compared to WT cells (Fig. S3). These results suggest that cells lacking EF-P are less susceptible to ciprofloxacin exposure, independent of homologous DNA repair activity.

**Figure 3.**
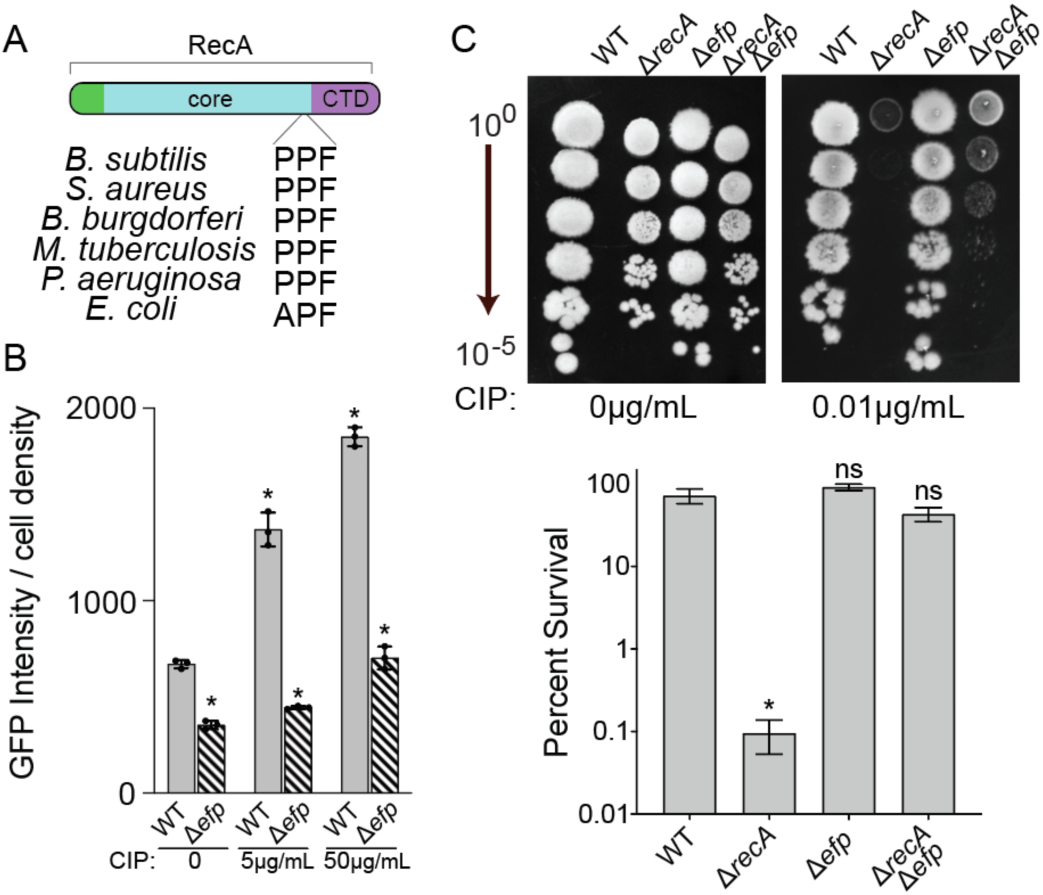
Loss of EF-P limits RecA production and confers survival of ciprofloxacin in the absence of RecA. A) Schematic of RecA and the location of the PPX conserved PPX site. B) Quantification of RecA-GFP production during exposure to ciprofloxacin. C) Representative plates and quantification of ciprofloxacin survival assays. Percent survival was calculated as colony forming units (CFU) on ciprofloxacin plates (50ng/mL) vs no drug controls. Data represent the mean ± SEM from biological replicate experiments (n=3). * Indicates p-value <0.05. ns indicates not statistically significant.

### RNA-seq reveals distinct transcriptional responses to ciprofloxacin in WT and Δ*efp* cells

To determine how EF-P loss affects the response to ciprofloxacin stress, we employed RNA-seq analysis to compare WT to a Δ*efp* mutant with and without ciprofloxacin treatment. Interestingly, most genes that are upregulated in response to ciprofloxacin are involved in the SOS response (Fig. 4A and Table S3). Cells lacking EF-P also upregulate genes involved in the SOS response pathway, albeit fewer genes are upregulated and the magnitude is less robust (Fig. 4B and Table S4). Genes involved in double-strand break repair and cell division checkpoints, including *recA*, *recN*, *yneA*, *dinB*, and the nucleotide excision repair factors *uvrA* and *uvrB*, were significantly upregulated in WT cells relative to cells lacking EF-P. In contrast, Δ*efp* cells showed muted induction of these SOS and repair genes. While several repair factors remained detectable, their expression levels were significantly lower compared to WT cells under identical drug exposure conditions.

**Figure 4.**
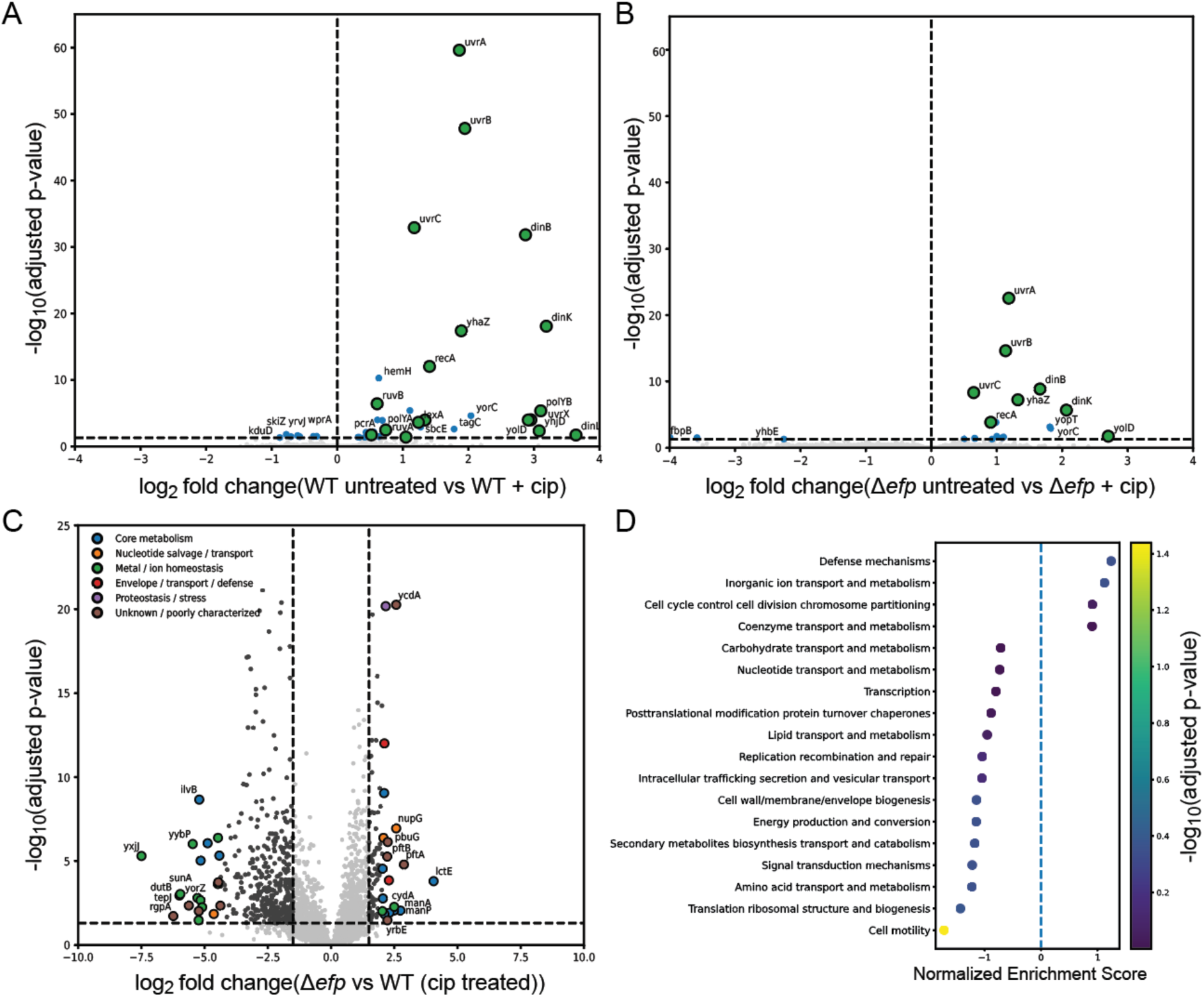
Transcriptomics analysis of WT and Δ*efp* cells during ciprofloxacin treatment. A) Analysis of WT (A) and Δ*efp* (B) transcriptome changes in untreated vs ciprofloxacin treated conditions. Green dots indicate SOS regulated genes. Blue dots indicate non-SOS significant hits. C) Transcriptomics analysis comparing *Δefp* and WT cells during ciprofloxacin treatment. The top 20 differentially expressed genes are labeled. D) Gene set enrichment analysis of the results in (C).

We next analyzed expression profiles from WT and Δ*efp* cells treated with ciprofloxacin (Fig. 4C and Table S5). In total, 613 genes were significantly differentially expressed. We performed gene set enrichment analysis (GSEA) to determine which functional categories were enriched in our dataset. Our analysis shows that genes involved in motility are significantly down regulated in cells lacking EF-P, consistent with previous reports (Fig. 4D) [24, 25]. Genes involved in cellular defense mechanisms were upregulated in cells lacking EF-P. The top two genes in this category were *bmrC* and *bmrD*, which encode subunits of a multidrug efflux pump (log_2_ fold-change of 1.93 and 2.11, respectively) (Table S5) [36]. We looked for other potential multidrug efflux pumps upregulated in our dataset and found that *bmrE* (log_2_ fold-change of 2.10) and *blt* (log_2_ fold-change of 1.10) were also differentially expressed. Although these two genes are not categorized under the defense mechanisms category, they have been implicated in antibiotic efflux [37, 38].

We observed similar patterns of differential expression when we compared WT and Δ*efp* cells in untreated conditions (Fig. S3 and Table S6). This is consistent with our observation that the primary differentially expressed genes in either WT or Δ*efp* cells in response to ciprofloxacin are SOS-regulated genes (Figs. 4A-B). Taken together, our results point to a model where the enhanced survival to ciprofloxacin of Δ*efp* cells results from transcriptional dysregulation, potentially mediated by the collective upregulation of efflux pumps, due to loss of EF-P and not a specific response to ciprofloxacin exposure.

## DISCUSSION

Antibiotic resistance continues to rise globally, underscoring the need to understand the full spectrum of cellular responses that modulate drug survival. While fluoroquinolone resistance is most often linked to mutations in DNA gyrase or Topoisomerase IV, our study reveals that *B. subtilis* regulates survival to ciprofloxacin treatment through EF-P-dependent processes [9]. Our study demonstrates that cells lacking EF-P show increased survival to sub-MIC concentrations of ciprofloxacin compared to WT cells. These results suggest that cells encode inherent mechanisms that can be leveraged to survive antibiotic exposure, allowing time to accumulate resistance mutations.

Studies in several experimental systems have shown that EF-P requires post-translational modification (PTM) to become functional [28, 29, 39, 40]. In *B. subtilis*, EF-P is post-translationally modified at lysine 32 with 5-aminopentanol, requiring multiple enzymes and modification intermediates [24, 30]. The regulatory function of this modification remains unclear. In previous studies, it has been shown that intermediate states of *B. subtilis* EF-P PTM may alter its function [30]. In our studies, we observed that loss of modification by YmfI phenocopies complete loss of EF-P. We found intermediate effects only in cells lacking GsaB. Future studies are needed to further understand the impact of intermediate PTM states on *B. subtilis* EF-P functionality. The regulation of EF-P PTM may provide a pathway for cells differentially regulate EF-P activity in response to environmental conditions, drastically altering cellular phenotypes, as has been shown the *Trypanosoma cruzi* EF-P homolog eIF5A [41].

In *B. subtilis*, ribosome profiling experiments have shown that EF-P regulated proteins are enriched for DNA replication and repair functions [25]. Our studies confirm that EF-P regulates the production of RecA, the enzyme that conducts homology search and catalyzes strand exchange during homologous recombination [42]. We show that in the absence of EF-P, RecA protein levels are reduced and unable to achieve full induction during exposure to DNA damaging agents. The results of our RNA-Seq experiment further confirm that cells lacking EF-P experience a muted SOS response compared to WT cells. This is counterintuitive to the increased survival to DNA damaging drugs that we observe. However, it has been shown that induction of the SOS regulon is not required to survive DNA damage in *B. subtilis* like it is in *E. coli* [35]. Furthermore, we show that in cells lacking RecA, loss of EF-P provides increased survival of ciprofloxacin treatment. Our RNA-Seq results show that several multidrug efflux pumps are upregulated due to loss of EF-P. Taken together, we prefer a model whereby loss of EF-P enhances survival to DNA damaging agents by limiting the intracellular concentration of the drugs, either through increased efflux or cell surface modifications that limit drug entry.

Our work demonstrates the role of EF-P in enhancing the susceptibility to a critical antibiotic. Further work is necessary to define the cumulative action of the pleiotropic effects of EF-P regulation. In *B. subtilis,* there are ∼780 polyproline motifs and EF-P regulates the production of 250 proteins during growth in rich media [25]. These factors make determining the exact contribution of EF-P regulation to a given phenotype difficult. While previous studies have shown that EF-P is required for homeostasis and stress responses during fast growth, our study is the first to observe a gain of function phenotype due to loss of EF-P [43, 44]. Understanding how EF-P regulates the susceptibility to fluoroquinolones will further inform the development of novel therapeutics by identifying critical pathways involved in drug efficacy.

## Supplementary Figures

**Supplementary Figure S1.**
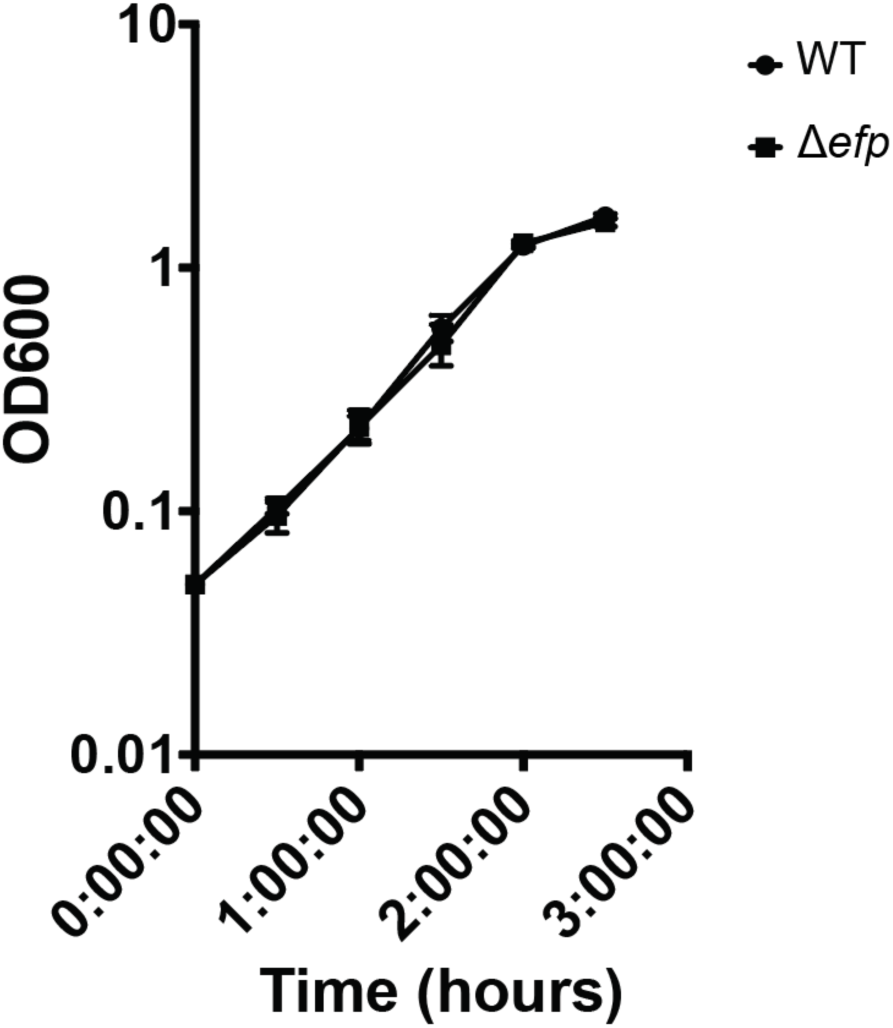
Grow curves of WT and Δ*efp* cells.

**Supplementary Figure S2.**
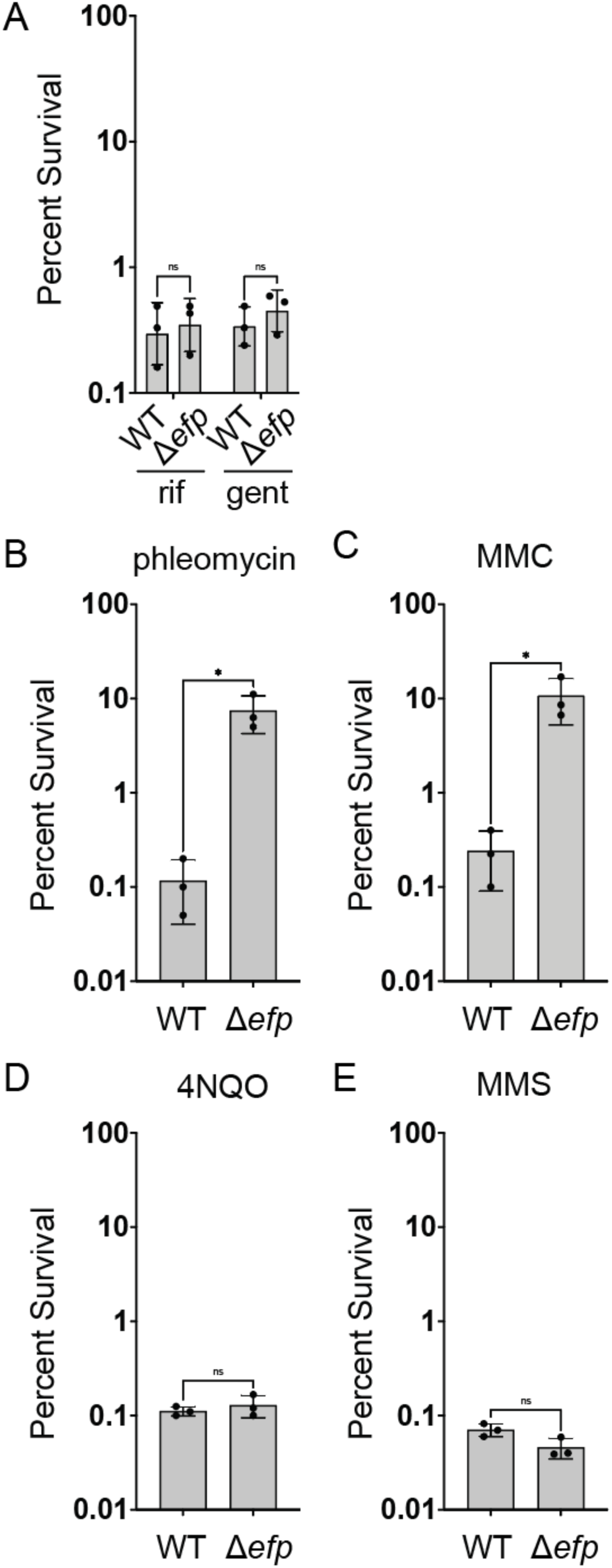
Cells lacking *efp* are less sensitive to potent double-strand break inducing drugs compared to WT, but not to other antibiotics.

**Supplementary Figure S3.**
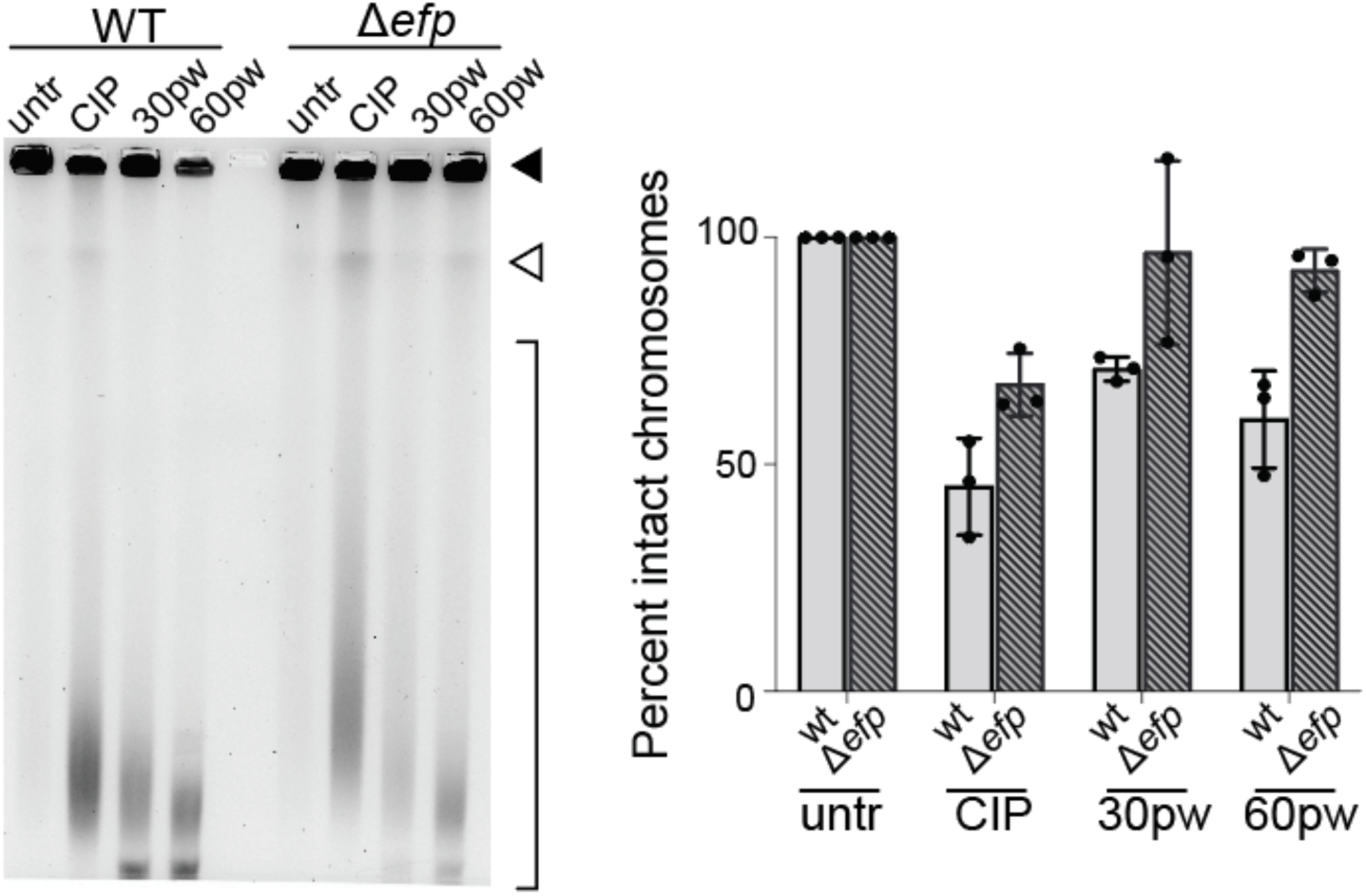
Cells lacking *efp* experience less double-strand breaks compared to WT cells after treatment with ciprofloxacin.

**Supplementary Figure S4.**
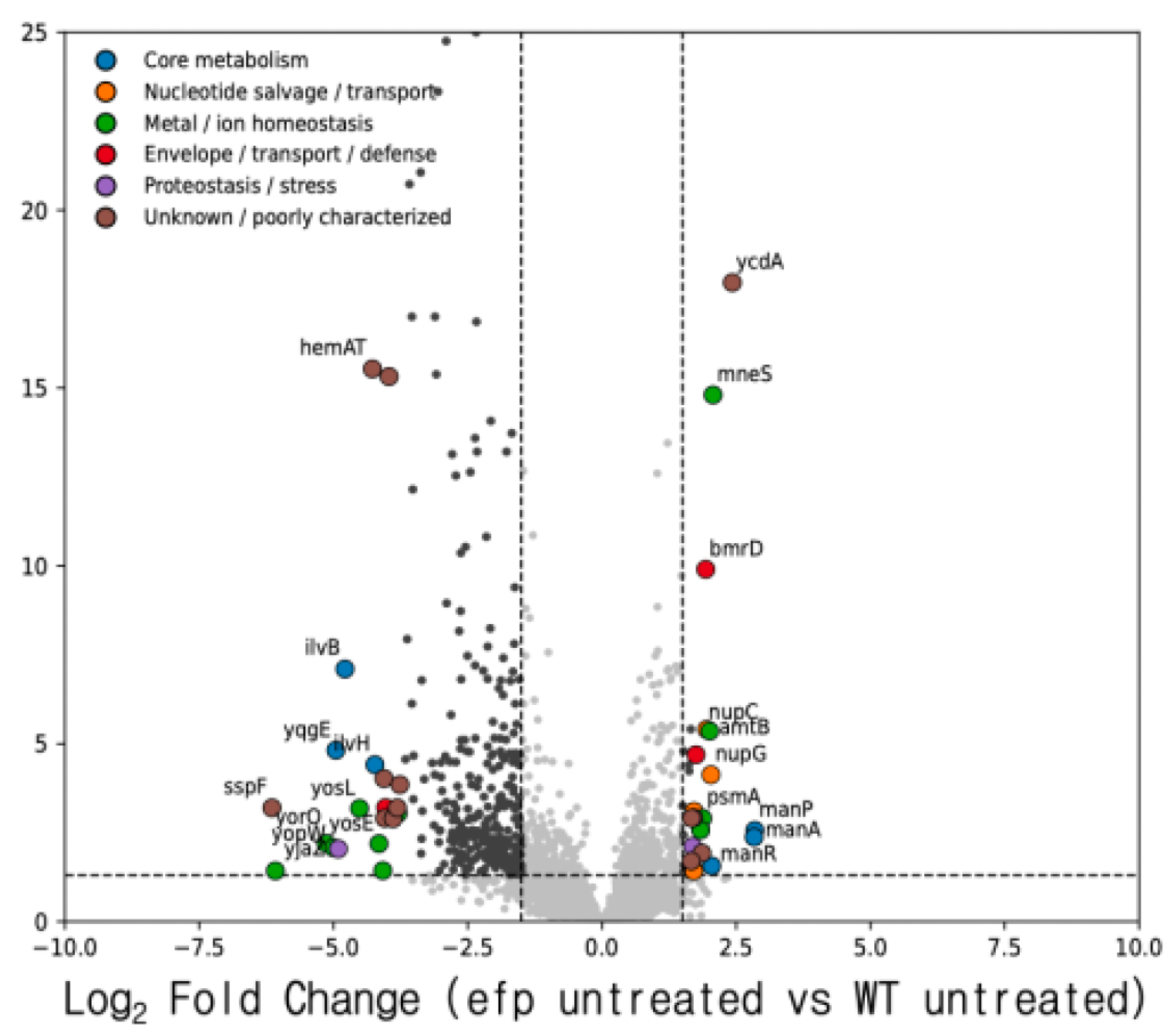
RNA-Seq analysis of WT and Δ*efp* cells in untreated conditions.

